# A novel ALDH1A1 inhibitor blocks platinum-induced senescence and stemness in ovarian cancer

**DOI:** 10.1101/2022.05.09.491218

**Authors:** Vaishnavi Muralikrishnan, Fang Fang, Tyler C. Given, Ram Podicheti, Mikhail Chchterbinine, Shruthi Sriramkumar, Heather M. O’Hagan, Thomas D. Hurley, Kenneth P. Nephew

## Abstract

Ovarian cancer is a deadly disease attributed to late-stage detection as well as recurrence and development of chemoresistance. Ovarian cancer stem cells (OCSCs) are hypothesized to be largely responsible for emergence of chemoresistant tumors. Although chemotherapy may initially succeed at decreasing the size and number of tumors, it leaves behind residual malignant OCSCs. In this study, we demonstrate that Aldehyde dehydrogenase 1A1 (ALDH1A1) is essential for the survival of OCSCs. We identified a first in class ALDH1A1 inhibitor, compound 974, and used 974 as a tool to decipher the mechanism of stemness regulation by ALDH1A1. Treatment of OCSCs with 974 significantly inhibited ALDH activity, expression of stemness genes, spheroid, and colony formation. In vivo limiting dilution assay demonstrated that 974 significantly inhibited CSC frequency. Transcriptomic sequencing of cells treated with 974 revealed significant downregulation of genes related to stemness and chemoresistance as well as senescence and senescence associated secretory phenotype (SASP). We confirmed that 974 inhibited senescence and stemness induced by platinum-based chemotherapy in functional assays. Overall, these data establish that ALDH1A1 is essential for OCSCs survival and ALDH1A1 inhibition suppresses chemotherapy induced senescence and stemness. Targeting ALDH1A1 using small molecule inhibitors in combination with chemotherapy therefore presents a promising strategy to prevent ovarian cancer recurrence and has potential for clinical translation.

## Introduction

Ovarian cancer is the most fatal gynecological malignancy[1]. In the US, ovarian cancer was the fifth leading cause of death among women and worldwide accounted for over 200,000 deaths in 2020[2]. High grade serous (HGS) is the most widely diagnosed subtype and accounts for 70-80% of ovarian cancer deaths [3]. Cytoreductive surgery and combination platinum-based chemotherapy have remained the mainstays of treatment. Although the majority of patients initially respond to chemotherapy, disease recurrence is common, and long-term survival in late-stage disease has improved little over the last four decades [4]. Mounting evidence shows that a small subpopulation of cells known as cancer stem cells (CSC) are associated with tumor relapse and chemoresistance in ovarian [5, 6] and other cancers [7]. Thus, it is essential to develop strategies to target CSC in conjunction with conventional therapies.

CSC are characterized by asymmetric division i.e., the ability to self-renew as well as differentiate into non-CSC, resistance to chemotherapy and radiation and the ability to survive without attachment. CSC are identified by biomarkers such as CD133[8], CD44/CD117[9], LGR5[10] or overexpression of aldehyde dehydrogenase (ALDH) enzymes[11]. ALDH1A1 is a member of the ALDH family and is highly expressed by stem cells in ovarian and other cancers[12]. Ovarian cancer cells with increased ALDH1A1 expression have higher self-renewal ability[13], and HGSOC patients with tumors expressing high ALDH1A1 have poor overall survival[11]. Although ALDH1A1 is a well-accepted marker for OCSC, the exact mechanism by which ALDH1A1 regulates stemness remains incompletely understood.

Stemness can be promoted by cellular senescence[14]. Senescence is a cellular state of irreversible growth arrest induced by oncogenic activation or DNA damaging therapies[15]. Senescent cells exhibit a complex secretome known as senescence associated secretory phenotype (SASP) consisting of cytokines, chemokines, and other growth factors. Senescence was initially thought to be tumor suppressive; however, recent evidence suggests that senescent cells have a pro-tumorigenic function[16]. In OC, platinum-based chemotherapy was shown to induce the CSC phenotype[17] and residual tumors after platinum treatment were enriched for ALDH+ cells[18]. Furthermore, platinum was shown to promote ovarian cancer stemness by paracrine signaling via SASP[19, 20], which could contribute to CSC enrichment.

To study the functional role of ALDH1A1 in OCSC, we identified a specific small molecule inhibitor, compound 974 (hereafter referred to as 974). This inhibitor acts as a unique tool to selectively block ALDH1A1 activity over other ALDH isoforms. We demonstrated that 974 inhibited stemness phenotypes in ovarian cancer cell lines expressing ALDH1A1 and in vivo limiting dilution analysis demonstrated an essential role for ALDH1A1 in CSC survival. Furthermore, transcriptomic sequencing of 974-treated HGSOC cells showed downregulation of pathways related to stemness and chemoresistance, including NFκB, IL6 signaling, xenobiotic metabolism, drug efflux and senescence, and 974 treatment blocked chemotherapy induced senescence and stemness. These results for the first time suggest a novel role for ALDH1A1 in maintenance of stemness via chemotherapy induced senescence in OC.

## Materials and Methods

### Chemical Reagents

Compounds purchased from ChemDiv Corporation (San Diego, CA) and ChemBridge Corporation (San Diego, CA) were >95% pure based on vendor specifications (NMR spectra for compounds can be found in the supplemental materials). Compound 974 was resynthesized in the IU Chemical Genomics Core facility, was greater than 99% pure by LC/MS and its structure validated by NMR.

### Protein Purification and Enzymatic Assays

Human ALDH1A1, ALDH1A2, and ALDH1A3 were prepared and purified as previously described[21-24]. Inhibition of ALDH activity by compounds and IC50 curves were determined by measuring the formation of NAD(P)H spectrophotometrically at 340 nm (molar extinction coefficient of 6200 M^-1^ cm^-1^) on the Beckman DU-640 as well as a Spectramax 340 PC spectrophotometer using purified recombinant enzyme. Reaction components for assays with ALDH1A enzymes consisted of 100-200 nM enzyme, 200 µM NAD+, 100 µM propionaldehyde, and 1% DMSO in 25 mM BES buffer, pH 7.5. All assays were performed at 25 °C and were initiated by addition of substrate after a 2 min incubation period. Purification of and reaction conditions for other ALDH isoenzymes were as described in[25]. IC50 curves were collected for compounds which substantially inhibited ALDH1A activity at 20 µM compound. Data were fit to the four parameter EC50 equation using SigmaPlot (v14), and the values represent the mean/SEM of three independent experiments (n = 3).

### X-ray Crystallography

All proteins used for crystallography were stored at -20 °C in 50 % (v/v) glycerol. Before use, proteins were dialyzed exhaustively against 10 mM ACES, 1 mM DTT, pH 6.6 buffer at 4°C. Crystals were grown using the sitting drop geometry at 20°C with crystallization solutions comprised of 100 mM BisTris pH 6.1-6.4, 9-11 % PEG3350 (Hampton Research, Catalog No. HR2-591), 200 mM NaCl, and 5 mM YbCl3. The complex with CM38 was made by soaking apo-enzyme crystals for 5 hours in the crystallization solution to which 500 µM compound in 2% DMSO (v/v) and 1 mM NAD+ had been added. The crystals were cryo-protected using 20 % ethylene glycol (v/v) in the same ligand soaking solution. Crystals were screened for diffraction on a Bruker X8 Prospector system. Diffracting crystals were stored in liquid nitrogen for transport to the synchrotron source. Diffraction data was collected at Beamline 19-ID of the Advanced Photon Source (Argonne National Laboratory, Chicago, IL). Data was integrated and scaled with the HKL3000 software suite. Rigid body, restrained TLS refinement and structure validation were performed using PHENIX (v1.17, 2-4). Modeling and visualization were performed using Coot (v0.8.9.2, 5) within PHENIX installation and PyMol v0.99 (DeLano Scientific LLC, San Francisco, CA).

### Cell culture

High grade serous ovarian cancer ovarian cancer cell lines OVCAR3, OVCAR5, OVSAHO and OVCAR8 were obtained from ATCC. OVCAR5 cell line was maintained in DMEM (Gibco, Catalog number: 11965092) with 10% FBS. All other cell lines were maintained in RPMI (Gibco, Catalog no. 11875135) with 10% FBS, 10ml of 100mM sodium pyruvate (Thermo Fisher, Catalog No. 11360070) and antibiotic-antimycotic (Thermo Fisher, Catalog No. 15240062). Cell lines were tested every 6 months for mycoplasma contamination using Mycoalert kit (Lonza, Catalog No. LT07-318).

### Flow cytometry

ALDH activity in live cells was measured by ALDEFLUOR Assay (Stem Cell Technologies, Catalog No. 01700) as per the manufacturer’s protocol. Percentage of ALDH+ cells was determined by LSRII flow cytometer (BD Biosciences), using 488 nm excitation and the signal was detected using the 530/30 filter. ALDH+ percentage gate was determined by sample specific negative control, Diethylamino benzaldehyde (DEAB)/ ALDH+ gate. CD133 was detected by flow cytometry using fluorescent-labelled antibody CD133/2-PE (Miltenyi Biotec, Catalog No. <u>130-120-145</u>) in LSRII flow cytometer using the filter 582/15. For each experiment, 10,000 events were analyzed. Flow cytometry data were collected using FACSDiva softw are (BD Biosciences) and analyzed using FlowJo software (FlowJo LLC).

### Quantitative PCR

RNA was isolated from cultured cells using RNeasy Mini Kit (Qiagen, Catalog No. 74104) following the manufacturer’s protocol. Nanodrop (ThermoFisher scientific) was used to determine RNA concentrations. qPCR was performed using Lightcycler 480 Kit (Roche Diagnostics, Catalog No. 04707516001) as described previously [17]. All gene expression data were normalized to human EEF1A1. Primer sequences are provided in Supplementary Table 2.

### Senescence Beta (β)-gal assay

Treated cells were stained for senescence-associated β-galactosidase activity according to manufacturer’s protocol (Cell Signaling Technology, Catalog No. 9860). The senescent cells were quantified by counting stained cells from five independent fields and percentage was calculated based on total number of cells in each field. Alternatively, percentage of senescence-associated β-galactosidase cells was determined by flow cytometry using SPiDER-β-Gal (Dojindo, Catalog No. SG-04) according to manufacturer’s protocol.

### RNA sequencing and bioinformatic analysis

OVCAR3 cells were treated with compound 974 (5 µM or DMSO for 48h in biological triplicates and total RNA was isolated using RNeasy Mini Kit (Qiagen, Catalog No. 74104) according to the manufacturer’s protocol. RNA-sequencing was performed essentially as we have described previously [26]. The RNA-seq results are available for download at Gene Expression Omnibus (GEO) data repository at the National Center for Biotechnology Information (NCBI) under the accession number GSE200641. See Supplementary Materials for a detailed description and bioinformatic analysis.

### Mouse xenograft experiment

All mouse experiments were performed according to ethical guidelines approved by the Institutional Animal Care and Use Committee of Indiana University (Bloomington, IN). For the limiting dilution analysis, 10^6^, 10^5^ or 10^6^ OVCAR3 cells of indicated conditions were mixed with Matrigel (Corning, Catalog No. CLS356234) at a 1:1 ratio and injected subcutaneously into right flanks of NOD SCID Gamma (NSG) mice. Tumor size was measured every week with a caliper and volume was calculated as ½*L*W^2^. At the end of the study, tumors were collected and dissociated using Tumor Dissociation Kit (Miltenyi Biotec, Catalog No.130-095-929) and a gentleMACS dissociator as per manufacturer’s protocol.

### Colony formation and tumorsphere assay

Cells at a 60-70% confluence in a 6-cm plate were treated for indicated times with the inhibitors. The cells were then collected by trypsinization and were plated as triplicates at a density of 500 cells/well in 24-well ultra-low adherent plates (Corning, Catalog No. 3473) with 1 mL of stem cell medium as described previously in [18] for spheroid formation assay or 6-well plates in 2-mL RPMI media with 10% FBS (for colony formation assay). Cells were allowed to grow for 7-14 days for spheroid formation or 5-7 days for colony formation. Spheroid size and morphology were assessed using a Zeiss Axiovert 40 inverted microscope with Axio-Vision software (Carl Zeiss Microimaging). Spheres larger than 10mm were counted under the microscope. Colonies were stained with 0.5% crystal violet and those with >50 cells were counted.

### MTT Cell proliferation assay

Cells were collected after inhibitor treatments by trypsinization and then were seeded at a density of 2000 cells per well in 96-well plates and 3-(4.5-dimethylthiazol-2-yl)-2.5-diphenyl tetrazolium bromide (MTT; Thermo Fisher Scientific, Catalog No. M6494) assay was performed at day as described previously[27]. IC_50_ values were calculated using Prism 7(GraphPad Software).

### Cell transfection and plasmids

100,000 OVCAR3 cells were transfected with shControl (Sigma-Aldrich, MISSION shRNA lentiviralSHC001V) or shALDH1A1(Sigma-Aldrich, MISSION shRNA lentiviralTRCN0000026415, TRCN0000026498,) as described in [18].

### Statistical analysis

All data are presented as mean values +/- SEM of at least three biological experiments unless otherwise indicated. Student t test was used to analyze the significant difference among different groups since the variation within the groups were similar. GraphPad Prism 7 software was used for data analysis and plotting.

### Data Availability

The data generated in this study are available within the article and its supplementary data files. The RNA-seq data generated in the study are publicly available in Gene Expression Omnibus (GEO) under the accession number GSE200641.

## Results

### Discovery of 974, a novel ALDH1A1 specific small molecule inhibitor

Compound 974 (974) is an ALDH1A1 specific small molecule inhibitor identified by screening compounds with high structural similarity to CM38, the lead compound identified from a high throughput screen [25]. CM38 showed good structural characteristics as a lead compound, with a low molecular weight of 294 kDa and an approximate ClogP of 2.8. To investigate the nature of the interactions that define inhibition in this series of compounds, we determined the structure of ALDH1A1 in a complex with both NAD and CM38 by X-ray crystallography to a resolution of 1.8 Å **(Supplementary Table S1**, PDB ID: 7UM9). The structure of CM38 bound to ALDH1A1 showed that CM38 bound within the substrate binding pocket of the enzyme **(Supplementary Fig. S1A)**. CM38 was then screened for ALDH inhibition using nine ALDH isoenzymes at 20 µM and showed excellent selectivity for ALDH1A1 over the other, highly similar isoenzymes in the ALDH subfamily **(Supplementary Fig. S1B)**. CM38 was found to be uncompetitive with respect to varied NAD+, which confirms that it does not bind the cofactor-binding site **(Supplementary Fig. S1C)**. There was no significant time-dependency in its ability to inhibit ALDH1A1, suggesting the interaction is non-covalent.

To avoid the potential off-target effects due to the structure of CM38, 974 was chosen amongst ALDH1A1 inhibitor with high structural similarity to CM38 from commercial sources (ChemDiv Corporation and ChemBridge Corporation). The structure of 974 is shown in **(Fig. 1A)**. Further details about discovery and characterization of 974 are described in supplementary methods. 974 is a highly potent inhibitor that blocks ALDH1A1 activity with an IC50 of 470nM **(Fig. 1B)**. 974 doses chosen for the rest of the study were lower than the IC50 doses for OVCAR3 and OVCAR5 cells **(Fig. 1C)**.

**Figure 1:**
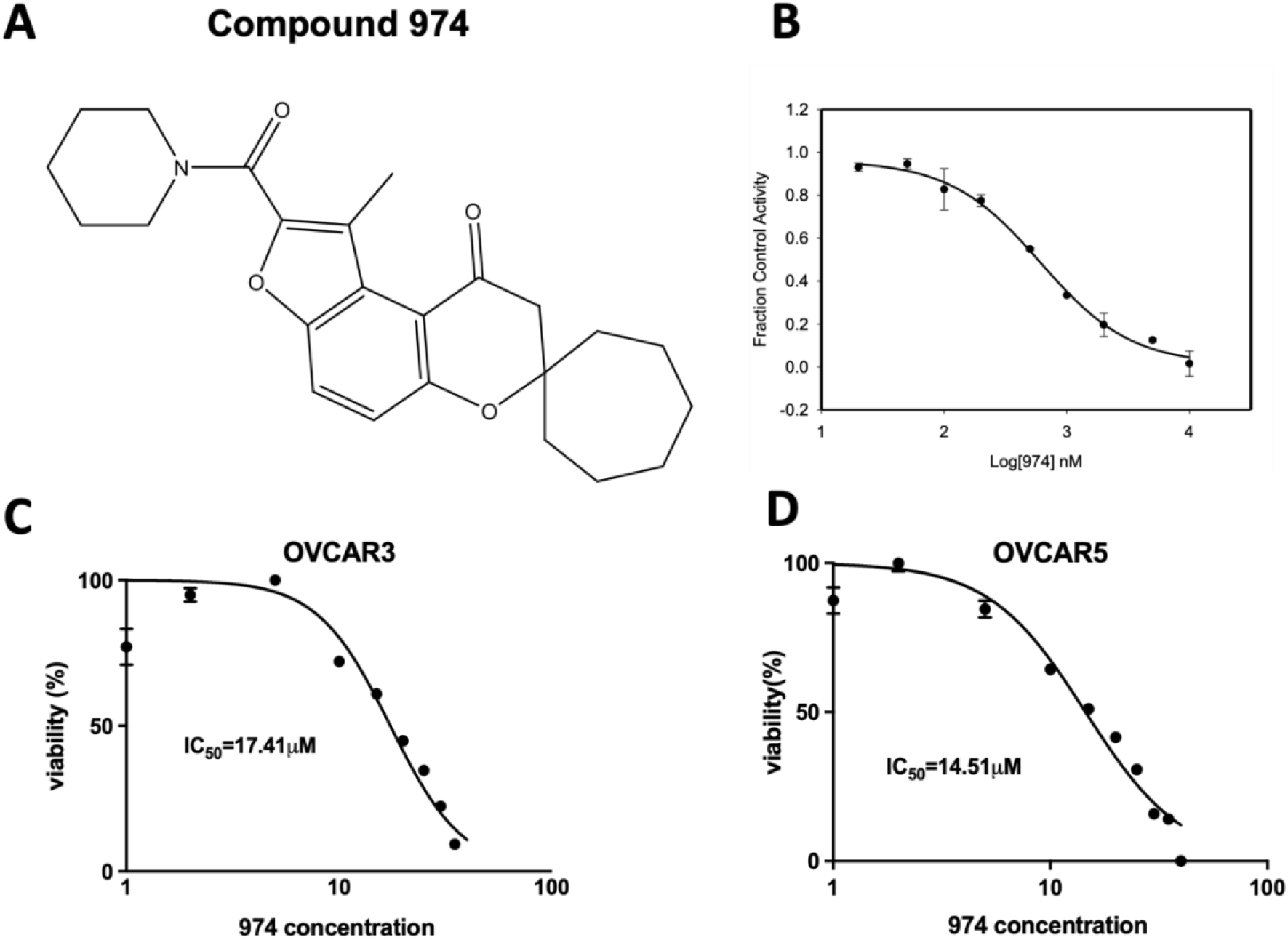
Compound 974: A novel ALDH1A1 inhibitor. **A**. Chemical structure of compound 974. **B**. EC50 curve for 974 binding with purified ALDH1A1. **C**. OVCAR3 (left) or OVCAR5 (right) were treated with increasing doses of 974 (0.5-100 µM) for 48 hours, and MTT assay was performed to measure viability. IC_50_ values were calculated using GraphPad Prism.

### ALDH1A1 inhibition suppresses stemness phenotypes ovarian cancer cells

To test the effect of 974 on cellular ALDH enzyme activity, we performed ALDEFLUOR assays in HGSOC cell lines. 974 significantly reduced the percentage of ALDH positive cells in OVCAR3 **(Fig. 2A)** and OVCAR5 **(Fig. 2B)**. The gating strategy for flow cytometry analysis for ALDEFLUOR assay is shown in **(Supplementary Fig. S2A)**. The dose of 974 was chosen based on a dose-response study **(Supplementary Fig. S2B)**. At the doses tested, 974 had no effect on the proliferation of ovarian cancer cells in monolayer **(Supplementary Fig. S2C)**. At the doses of 974 selected for further study, the compound did not induce apoptosis, indicated by PI/Annexin V staining **(Supplementary Fig. S2D)**.

**Figure 2.**
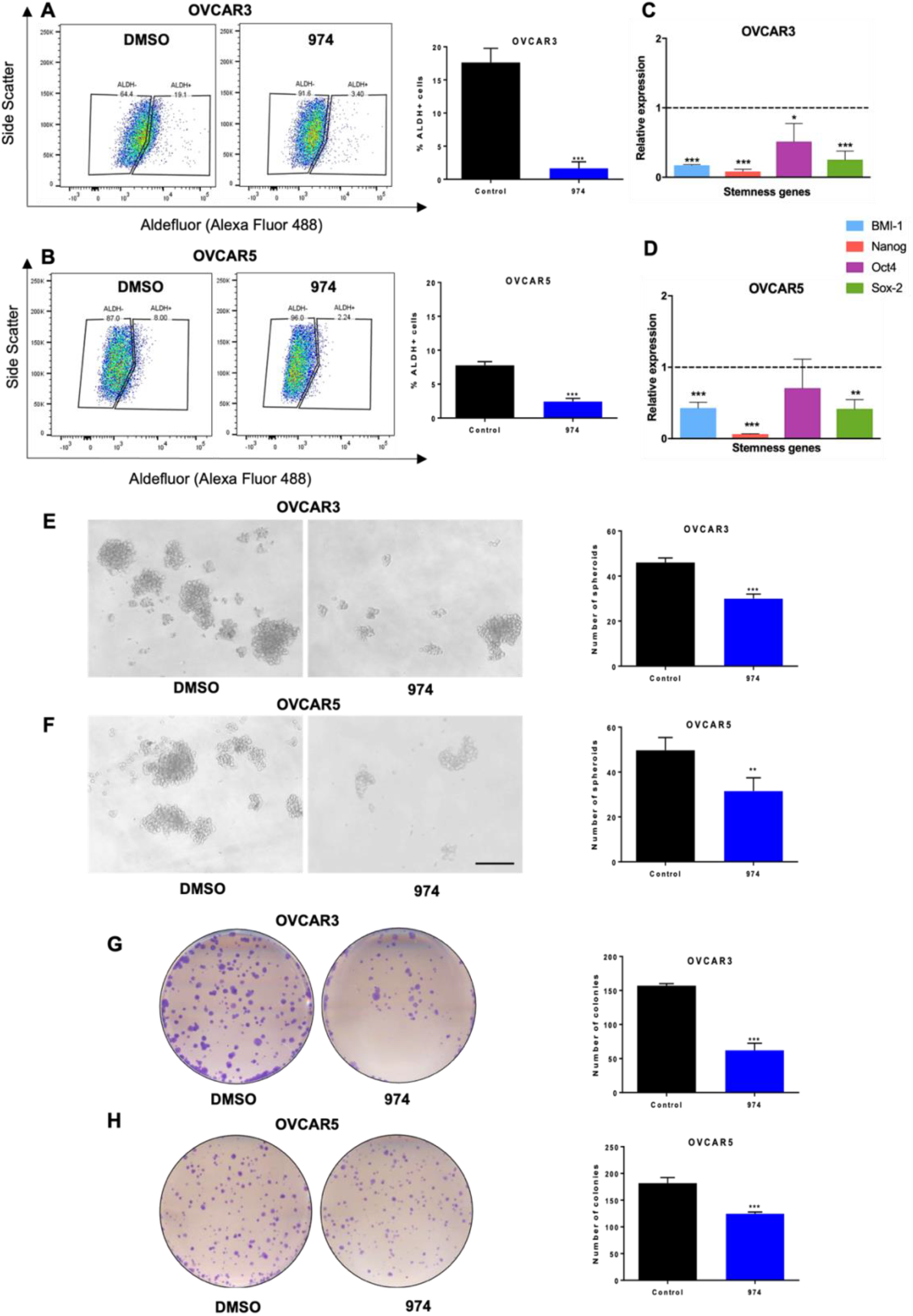
ALDH1A1 inhibition suppresses ovarian cancer stemness phenotypes in vitro. **A**. OVCAR3 or **B**. OVCAR5 cells were treated with compound 974 (5µM for 48h) or DMSO and the percentage of ALDH+ cells were measured by ALDEFLUOR assay using flow cytometry (left) and the results were quantified (right). **C**. OVCAR3 or **D**. OVCAR5 cells were treated as in A and expression of stemness genes were measured by qPCR. **E**. OVCAR3 or **F**. OVCAR5 cells were treated as in A. 500 cells/well were replated in 24-well low-adhesion conditions after treatment. Representative images of spheroid formation after 14 days (left) and quantification (right). **G**. OVCAR3 or **H**. OVCAR5 were treated as in A. 500 cells/well were replated in 6 well plates after treatment. Colonies were stained with 0.05% crystal violet and counted. Representative images of colony formation (left) and quantification (right). Error bars represent SEM; n =3 independent experiments of triplicate assays. Data are presented as mean ± SEM with p < 0.05 (*), p < 0.01 (**), and p < 0.005 (***). Scale bar, 100µm.

Numerous genes associated with stemness have been reported to be characteristics of OCSCs [28, 29]. 974 significantly decreased expression of well-known stemness genes Bmi-1, Nanog, Oct4, and Sox2 in OVCAR3 **(Fig. 2C)** and OVCAR5 **(Fig. 2D)**. To measure the self-renewal ability of the CSC subpopulations, the spheroid formation assay was used. Spheroid formation ability of OVCAR3 **(Fig. 2E)** and OVCAR5 **(Fig. 2F)** was significantly inhibited by 974 treatment, and clonogenic survival of both cell lines was also significantly inhibited by treatment with 974 **(Fig 2. G, H)**. To determine if the effects of 974 were specific to ALDH1A1-mediated stemness, we used OVCAR8 cells, which do not have detectable ALDH activity but do have a CD133+ stem cell population (**Supplementary Fig. S3A)**. 974 treatment did not alter clonogenic growth or spheroid formation in OVCAR8 (**Supplementary Fig. S3B-D)**. Treatment with 974 also did not alter the percentage of CD133 cells in OVCAR8 **(Supplementary Fig. S3A)**.

To determine if genetically reducing ALDH1A1 levels had similar effects on stemness properties as drug treatment, we developed stable 2 independent shRNA mediated ALDH1A1 knockdown (shALDH1A1_1 and shALDH1A1_2) and scrambled control (shControl) OVCAR3 cell lines **(Supplementary Fig. S4A)**. ALDH1A1 knockdown significantly decreased the percent ALDH+ cells, spheroid, and colony formation compared to shControl **(Supplementary Fig. S4B-D)**, similar to what was observed by 974 treatment. To test the specificity of 974 to ALDH1A1 in cells, shALDH1A1 and shControl cells were treated with 974 and ALDEFLUOR assay was performed. 974 did not further reduce the percentage of ALDH+ cells in shALDH1A1 cells **(Supplementary Fig. S5)**.

### ALDH1A1 inhibition suppresses ovarian cancer stemness in vivo

To test whether 974 treatment blocks tumor initiation *in vivo*, a limiting dilution analysis (LDA) was performed. OVCAR3 cells were pretreated with 974 (5µM) or DMSO for 48 hours and 1 million, 100,000 or 10,000 treated cells were injected subcutaneously (s.c.) into NSG mice and tumor formation was monitored **(Fig. 3A)**. Treatment with 974 significantly reduced CSC frequency in mice **(Fig. 3B)**. Complementary to the study using 974 and to examine the requirement for ALDH1A1 in this context, shALDH1A1 or shControl cells (1 million, 100,000 or 10,000) were injected s.c in NSG mice. The results of the LDA demonstrated a significant reduction in CSC frequency **(Fig. 3C)**. The log fraction plot was generated using the LDA software for the 974(or DMSO) study as well as the shALDH1A1(or shControl) study **(Fig. 3D, E)**. The slope of the solid line represents the log-active cell (CSC) fraction. The 95% confidence interval is shown by the dotted lines. At the end of the study, tumors were collected and analyzed for percentage of ALDH+ cells by ALDEFLUOR assay. The tumors from mice injected with shALDH1A1 cells had significantly lower percentage of ALDH+ cells compared to the tumors from shControl injected mice **(Fig. 3F)**. Collectively, these data demonstrate that ALDH1A1 is essential for maintenance of stemness in ovarian cancer cells and 974 significantly inhibits stemness phenotypes.

**Figure 3.**
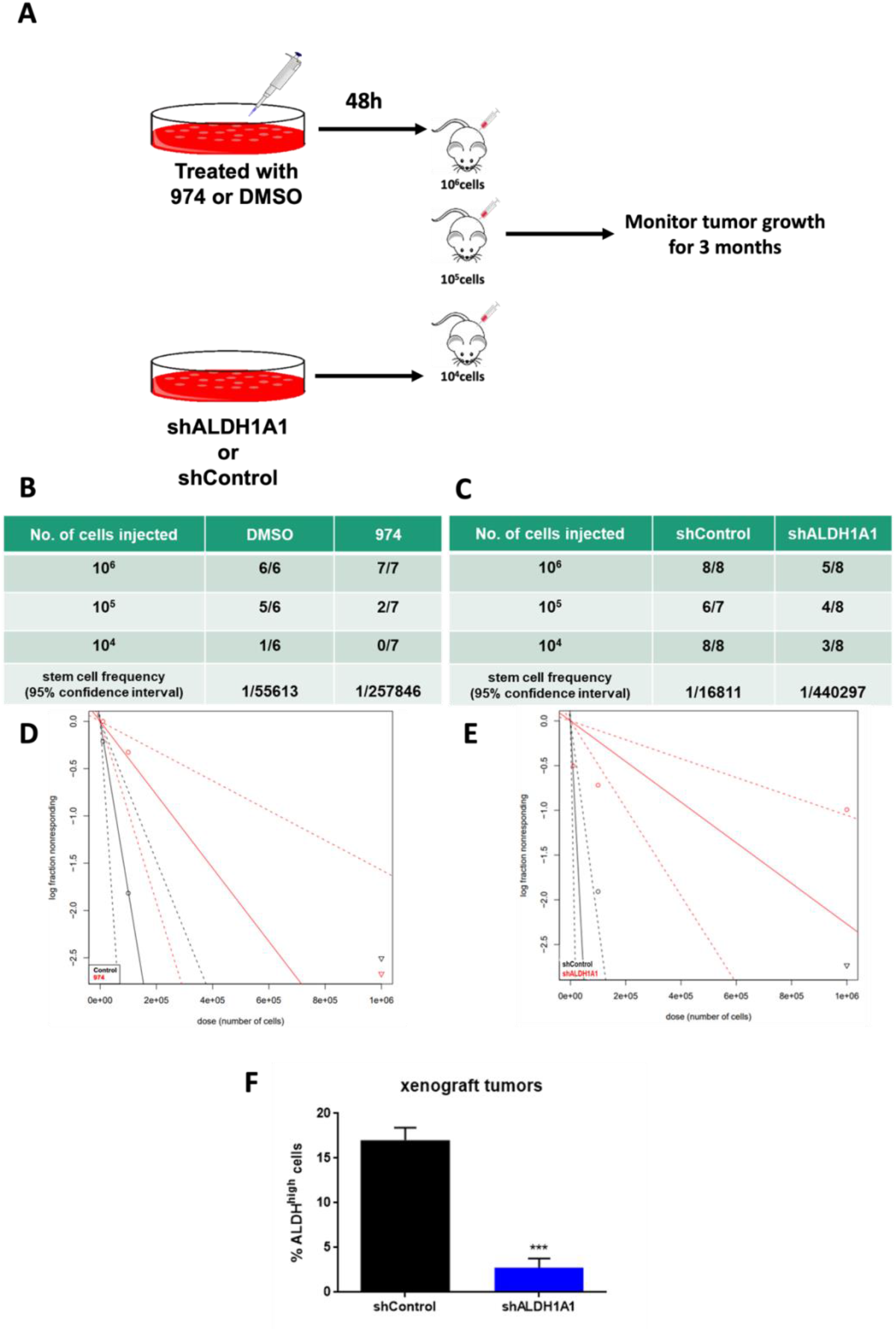
ALDH1A1 inhibition suppresses ovarian cancer stemness in vivo. **A**. Schematic representing study design. 10^6^, 10^5^ or 10^4^ OVCAR3 cells treated with compound 974 (5µM for 48h) or DMSO OR shALDH1A1 or shControl cells were injected into NSG mice subcutaneously and tumor formation was monitored. Numbers of mice with tumors over the total numbers of mice in the group and CSC frequency calculated by ELDA software (https://bioinf.wehi.edu.au/software/elda/) **B**. For DMSO or 974 treatment **C**. For shControl or shALDH1A1. Log-fraction plot of limiting dilution analysis for stem cell frequency generated by extreme limiting dilution analysis in **D**. compound 974 treatment vs DMSO or **E**. shALDH1A1 or shControl. **F**. Percentage of ALDH+ cells in the dissociated tumors from shALDH1A1 study in C was measured by ALDEFLUOR assay. Error bars represent SEM; n =3 independent tumor samples. Data are presented as mean ± SEM with p < 0.005 (***).

### ALDH1A1 inhibition downregulates key stemness and chemoresistance pathways

To determine the effect of ALDH1A1 inhibition by 974 on gene expression, transcriptomic analysis of 974 or DMSO treated cells was carried out using RNA-seq and bioinformatic analysis. The volcano plot shows that 1630 genes were downregulated, and 1140 genes upregulated by ALDH1A1 inhibition **(Fig. 4A**, 974 vs. DMSO treated samples; FDR<0.05). Genes significantly downregulated by ALDH1A1 inhibition included stem cell markers (CD44, FZD7, SOX9) and genes involved in chemoresistance (ABCB1, NFκB) in ovarian cancer [9, 30-32] **(Fig. 4B)**. In addition, 974 treatment significantly downregulated senescence biomarkers p21(CDKN1A) and p15^INK4b^ (CDKN2B) and genes associated with the senescence associated secretory phenotype (SASP) including IL6, IL8, CXCL1, CXCL3 **(Fig. 4B)**. Ingenuity Pathway Analysis (IPA) and Gene Ontology (GO) annotations of the differentially expressed genes demonstrated that 974 treatment inhibited a number of key biological processes associated with tumor initiation and stem cells, including growth of solid tumor, inflammatory response, cells movement of cancer cells, development of epithelial tissues and drug resistance of tumor cells (red and green colors represent upregulated or downregulated genes respectively), supporting the role of ALDH1A1 in modulating OCSC biology **(Fig. 4C)**. IPA of genes significantly decreased by 974 treatment revealed altered xenobiotic metabolism signaling, cancer drug efflux, IL6 and NFκB signaling (**Fig. 4D, Supplementary Table S4)**, all of which have been reported to play a role in stemness and chemoresistance[18, 33, 34]; furthermore, the cellular senescence pathway was significantly downregulated by 974 treatment **(Fig. 4D)**. Collectively, these results demonstrated that ALDH1A1 inhibition led to reduction in gene expression in stemness and chemoresistance related pathways in OC.

**Figure 4.**
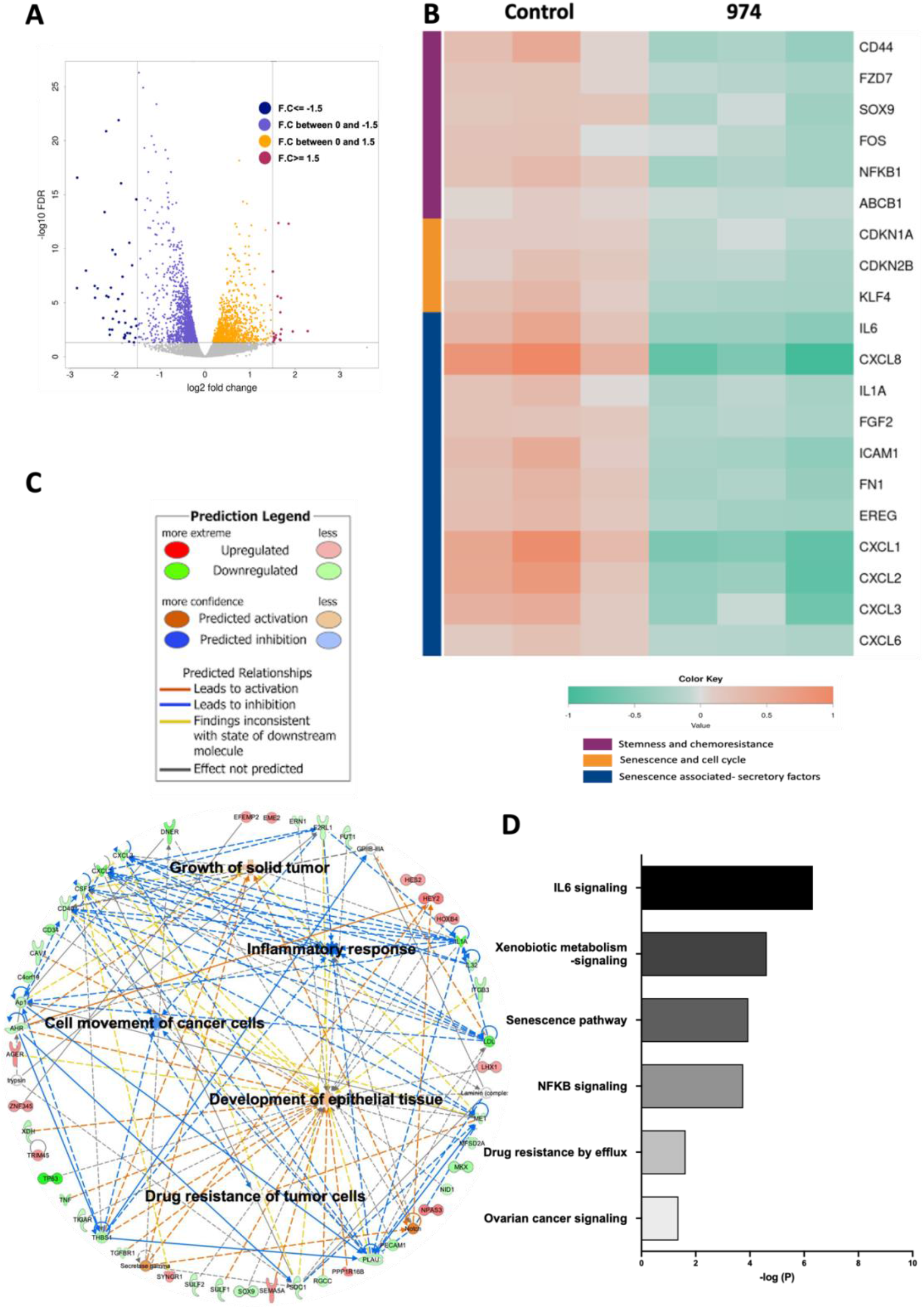
ALDH1A1 inhibition suppresses pathways involved in chemoresistance and stemness. RNA-seq was performed on OVCAR3 cells treated with compound 974 (5µM for 48h) or DMSO (n=3). **A**. Volcano plot of genes up and downregulated by ALDH1A1 inhibition. **B**. Heatmap of selected genes significantly downregulated by compound 974 (FDR < 0.05). **C**. Networks of biological processes constructed using significantly altered genes (FDR < 0.05) between OVCAR3 cells treated with 974 or DMSO. **D**. Canonical pathways related to stemness and chemoresistance identified by Ingenuity pathway analysis (IPA) using genes significantly altered by 974 treatment (FDR < 0.05, Fold change>|1.5|).

### Inhibition of ALDH1A1 suppresses chemotherapy induced senescence and stemness

Senescence is the cellular state characterized by proliferative arrest, resistance to apoptosis and altered expression of genes encoding cytokines and other growth factors, commonly known as SASP[35]. SASP has pro-tumorigenic paracrine effects, and emerging evidence supports the role of the SASP in the induction of cancer stemness and relapse [14, 20]. Because platinum-based chemotherapy has been shown to enhance SASP and subsequently stemness in ovarian cancer [20], we investigated the effect of ALDH1A1 inhibition on senescence in cisplatin (CDDP)-treated cells. Treatment with 974 suppressed CDDP-induced senescence associated beta galactosidase (SA-β gal) staining in OVCAR3 **(Fig. 5A, B)**. 974 treatment also significantly reduced basal and CDDP-induced expression of senescence marker p21 (CDKN1A) in OVCAR3 **(Fig. 5C)** and SASP gene expression in OVCAR3 and OVCAR5 **(Fig. 5C; Supplementary Fig. S6A)**. Additionally, 974 significantly inhibited SASP gene expression in CDDP-resistant OVCAR5 cells with high ALDH activity, developed by repeated cycles of exposure to CDDP **(Supplementary Fig. S6B-D, Supplementary Methods)**. The effect of ALDH1A1 inhibition on senescence phenotype was validated by using the shALDH1A1 cells. ALDH1A1 knockdown significantly suppressed basal and CDDP-induced senescence measured as percentage of β-gal positive cells **(Fig. 5D)**. ALDH1A1 knockdown suppressed the basal and CDDP-induced expression of p21 and SASP genes, including IL6, IL8, CXCL1 and CXCL3 **(Fig. 5E)**.

**Figure 5.**
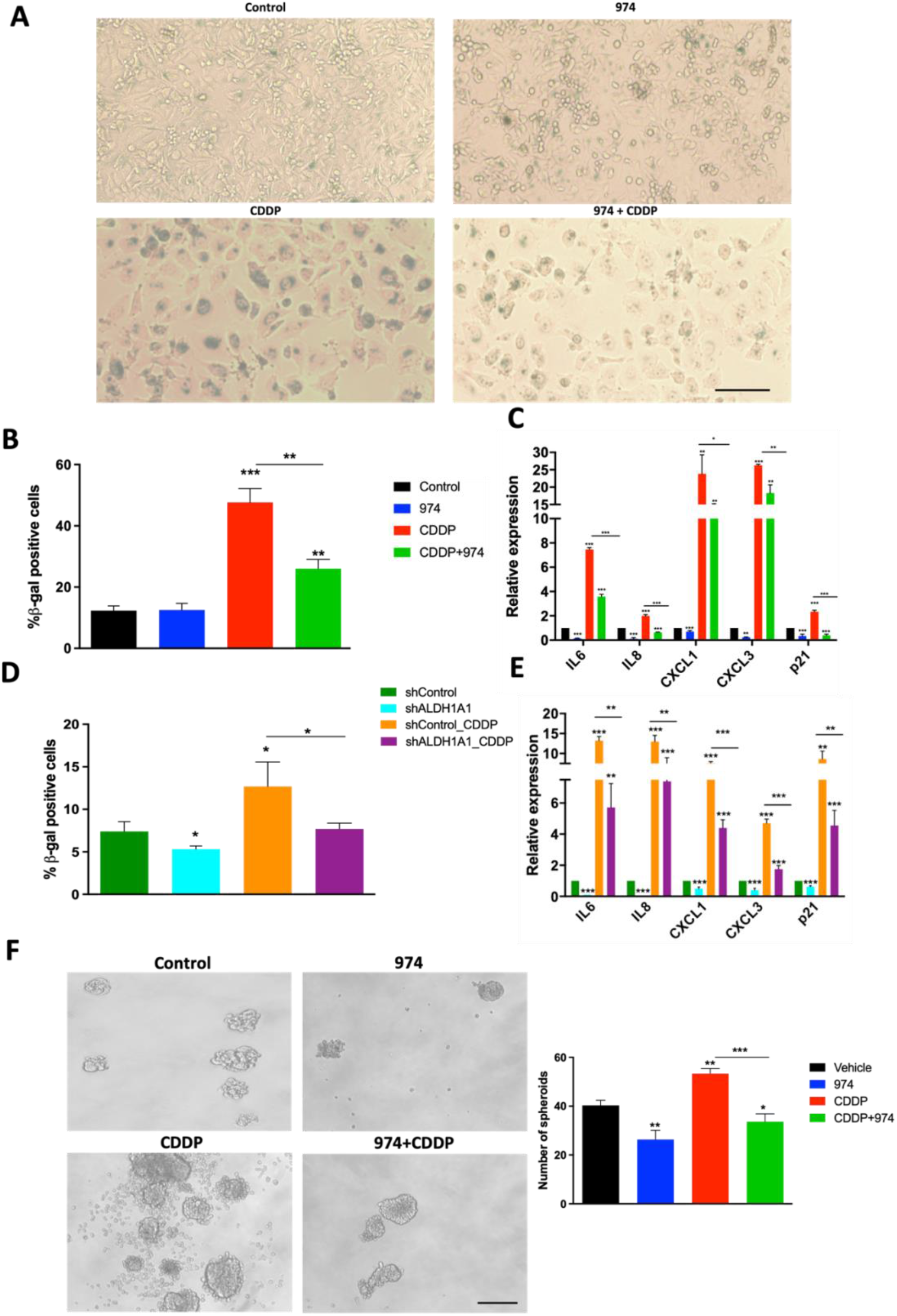
ALDH1A1 inhibition suppresses chemotherapy induced senescence and stemness in HGSOC cells. **A**. Senescence associated (SA) Beta-gal assay was performed on OVCAR3 cells treated with DMSO, compound 974 (5µM for 48h), cisplatin (CDDP) (15µM for 16h) or both. Representative images at 10X magnification. **B**. Quantification of SA-Beta-gal assay represents percentage of senescent cells averaged from 5 different fields in each condition. **C**. Expression of SASP genes and p21 (CIP1/WAF1) was examined by qPCR in OVCAR3 cells treated as in A. **D**. Percentage of SA-Beta gal positive cells in shControl or shALDH1A1 cells treated with NaCl (vehicle) or CDDP (15µM for 16h) was measured by flow cytometry using Spider beta gal reagent. **E**. Expression of p21 and SASP genes was examined by qPCR in shControl or shALDH1A1 cells treated with NaCl or CDDP (15µM for 16h). **F**. OVCAR3 cells treated with DMSO, compound 974 (5µM for 48h), cisplatin (CDDP) (7.5µM; _½_ IC50 for 3h) or both were plated in low-attachment conditions at a density of 500 cells/well. Representative spheroid images captured on Day 7 (left). Quantification of spheroids (right). Error bars represent SEM; n =3 independent experiments of triplicate assays. Data are presented as mean ± SEM with p < 0.05 (*), p < 0.01 (**), and p < 0.005 (***). Scale bar, 100µm.

CDDP induces stemness in ovarian cancer cells and treatment with 974 abrogated CDDP-induced stemness phenotype in spheroid assays **(Fig. 5F)**, suggesting a link between senescence and stemness. To confirm that blocking senescence reduced stemness, ovarian cancer cells were treated with ABT-263, a senolytic agent (Navitoclax; 2µM for 24h). ABT-263 treatment reduced basal and CDDP-induced spheroid numbers **(Supplementary Fig. S6C)**.

## Discussion

Ovarian cancer is a deadly disease attributed to late-stage detection as well as relapse and development of chemoresistance. Strategies to overcome chemoresistance are needed to achieve better prognosis in ovarian cancer patients. OCSCs have been shown to cause chemoresistance[17], thus targeting this population in conjunction with conventional chemotherapy could be an effective strategy in preventing relapse. This study describes the discovery and characterization of compound 974, a novel small molecule inhibitor selective to ALDH1A1 over other ALDH isoforms. We show that 974 inhibits stemness phenotypes in HGSOC cell lines, blocks expression of putative stemness genes and pathways, reduces OCSC frequency and delays tumor initiation *in vivo*. Importantly, inhibition of ALDH1A1 by 974 suppresses platinum-based chemotherapy induced senescence and stemness and to our knowledge this is the first report that ALDH1A1 regulates senescence-mediated stemness. Overall, our findings support the use of small molecule inhibitors of ALDH1A1 as a promising therapeutic approach to target OCSC and prevent chemoresistance.

ALDH1A1 is a robust marker for CSCs in ovarian and other cancers [12, 36] and ALDH1A1 expression in patient tumors predicts poor prognosis[11, 37]. Several inhibitors have been designed to target CSCs by selective inhibition of ALDH1A1 or by a pan-ALDH1A inhibition approach. Pan-ALDH1A inhibitor 673A causes cell death by necroptosis in OCSCs, reduces tumor initiation and is highly synergistic with chemotherapy[38]. Pan-ALDH1A inhibitor has the advantage of overcoming resistance that arises from compensation due to different ALDH isoforms. However, targeting ALDH1A1 selectively could have a unique advantage from safety perspective since ALDH1A1 has been shown to be dispensable for stem cell function in mice [39]. Moreover, ALDH1A2 expression is essential for dendritic cell differentiation in bone marrow microenvironment [40].

By targeting a novel scaffold in ALDH1A1, 974 acts an effective tool for understanding ALDH1A1 function. We show that 974 is specific to ALDH1A1 by demonstrating no change in stemness when treating a low ALDH1A1 expression cell line with 974 (OVCAR8 cells**; Supplementary Fig. S3 A-D)**. Moreover, when ALDH1A1 is biologically inhibited using a knockdown, 974 can no longer inhibit ALDH activity **(Supplementary Fig. S5)**. The effects are in line with the published reports of other ALDH1A1 specific inhibitors CM37[13] and NCT-501[41]. The data on ALDH1A1 inhibition using CM37 and NCT-501 support our findings that ALDH1A1 is essential to maintain CSC phenotypes. CM37 inhibited spheroid formation and expression of stemness genes such as Sox2, Nanog, Oct4 as well as p21 similar to 974[13]. NCT-501 inhibits ALDH activity and attenuates de-differentiation of non-CSC into CSCs in ovarian cancer cells[41]. Indirect targeting of ALDH1A1 includes therapies that target upstream ALDH1A1 regulators of such as BRD4[42] or HOTAIR[43] thereby inhibiting the expression of ALDH1A1 which leads to inhibition of stemness. Although 974 is not yet formulated as an in vivo therapeutic, we provide compelling evidence using ovarian cancer cells treated with 974 in vitro for LDA as support for future rational chemistry design strategies to improve 974 bioavailability and targeting CSC in vivo. Our ongoing efforts aim at targeting the “arms” that extend from the central scaffold to improve the metabolic stability and modify the lipophilicity of the compound.

ALDH1A1 is a ubiquitous enzyme with several cellular functions such as conversion of aldehydes into carboxylic acids, scavenging ROS, altering signaling through modulation of retinoic acid pathway[12]. Thus, how ALDH1A1 regulates cancer stemness could involve more than one mechanism. Through transcriptomic analysis, our study reveals a previously unknown mechanism of stemness regulation by ALDH1A1 via the senescence pathway. Senescence, initially thought to be a tumor suppressive mechanism, has recently been shown to promote stemness in ovarian[20] and other cancers[16]. Senescent cells exhibit a complex secretome known as SASP that promotes stemness via paracrine signaling [20]. Specifically, IL-6 signaling axis was shown to be upregulated by neo-adjuvant chemotherapy[44]. We demonstrate that ALDH1A1 inhibition blocks the senescence and SASP induced by cisplatin treatment (Figure 5), possibly by suppressing the NFκB pathway (Figure 4D). NFκB regulates several of the SASP factors[36], and inhibiting NFκB attenuates stemness in OC[45]. Further investigation is required to elucidate the exact mechanism by which ALDH1A1 regulates cisplatin-induced stemness.

In conclusion, using 974 as a tool, we have demonstrated the functional significance of ALDHA1 in maintaining ovarian cancer stemness *in vitro* and *in vivo* models. To our knowledge, this is the first study that demonstrates that ALDH1A1 is involved in regulation of senescence and SASP. ALDH1A1 regulation of senescence could be significant because standard of care treatment for ovarian cancer includes platinum-based chemotherapy, and carboplatin has been shown to induce senescent cells in ovarian cancer tumors[46, 47]. We show that a new isoform-specific ALDH1A1 inhibitor suppresses chemotherapy-induced senescence as well as stemness. Targeting ALDH1A1 in combination with chemotherapy could block senescence and inhibit CSC enrichment to overcome resistance and improve outcomes for ovarian cancer patients.

## Conclusion

Here we describe the, a new specific and potent small molecule inhibitor for ALDH1A1 in ovarian cancer models enriched in cells with stemness characteristics. Together with siRNA-mediated knockdown of the enzyme, the results obtained using CM37 provide proof of principle supporting the role of ALDH1 in cancer stemness. Our data demonstrate that by fine tuning the levels of intracellular oxidative stress, ALDH1A1, protects cancer cells from DNA damage, enhancing spheroid proliferation and tumorigenicity.

## Acknowledgements

We thank Christiane Hassel (Flow Cytometry Core Facility, Indiana University, Bloomington, IN) for technical assistance with flow cytometry. We would like to thank the Center for Genomics and Bioinformatics at Indiana University, Bloomington, for their assistance with RNA-seq experiments, especially Jie Huang for library construction and sequencing. This project was funded with support from the Indiana Clinical and Translational Sciences Institute funded, in part by Grant Number UL1TR002529 from the National Institutes of Health, National Center for Advancing Translational Sciences, Clinical and Translational Sciences Award; and through the IU Simon Comprehensive Cancer Center P30 Support Grant (P30CA082709-20).

## References

1. Torre, L.A., et al., Ovarian cancer statistics, 2018. CA Cancer J Clin, 2018. 68(4): p. 284–296.

2. Sung, H., et al., Global Cancer Statistics 2020: GLOBOCAN Estimates of Incidence and Mortality Worldwide for 36 Cancers in 185 Countries. CA Cancer J Clin, 2021. 71(3): p. 209–249.

3. Kurman, R.J. and M. Shih Ie, The Dualistic Model of Ovarian Carcinogenesis: Revisited, Revised, and Expanded. Am J Pathol, 2016. 186(4): p. 733–47.

4. Davis, A., A.V. Tinker, and M. Friedlander, “Platinum resistant” ovarian cancer: what is it, who to treat and how to measure benefit? Gynecol Oncol, 2014. 133(3): p. 624–31.

5. Steg, A.D., et al., Stem cell pathways contribute to clinical chemoresistance in ovarian cancer. Clin Cancer Res, 2012. 18(3): p. 869–81.

6. Al-Alem, L.F., et al., Ovarian cancer stem cells: What progress have we made? Int J Biochem Cell Biol, 2019. 107: p. 92–103.

7. Dean, M., T. Fojo, and S. Bates, Tumour stem cells and drug resistance. Nat Rev Cancer, 2005. 5(4): p. 275–84.

8. Baba, T., et al., Epigenetic regulation of CD133 and tumorigenicity of CD133+ ovarian cancer cells. Oncogene, 2009. 28(2): p. 209–18.

9. Zhang, S., et al., Identification and characterization of ovarian cancer-initiating cells from primary human tumors. Cancer Res, 2008. 68(11): p. 4311–20.

10. Yu, S., et al., Therapeutic Targeting of Tumor Cells Rich in LGR Stem Cell Receptors. Bioconjug Chem, 2021. 32(2): p. 376–384.

11. Landen, C.N., Jr., et al., Targeting aldehyde dehydrogenase cancer stem cells in ovarian cancer. Mol Cancer Ther, 2010. 9(12): p. 3186–99.

12. Muralikrishnan, V., T.D. Hurley, and K.P. Nephew, Targeting Aldehyde Dehydrogenases to Eliminate Cancer Stem Cells in Gynecologic Malignancies. Cancers (Basel), 2020. 12(4).

13. Nwani, N.G., et al., A Novel ALDH1A1 Inhibitor Targets Cells with Stem Cell Characteristics in Ovarian Cancer. Cancers (Basel), 2019. 11(4).

14. Milanovic, M., Y. Yu, and C.A. Schmitt, The Senescence-Stemness Alliance - A Cancer-Hijacked Regeneration Principle. Trends Cell Biol, 2018. 28(12): p. 1049–1061.

15. Herranz, N. and J. Gil, Mechanisms and functions of cellular senescence. J Clin Invest, 2018. 128(4): p. 1238–1246.

16. Milanovic, M., et al., Senescence-associated reprogramming promotes cancer stemness. Nature, 2018. 553(7686): p. 96–100.

17. Wang, Y., et al., Epigenetic targeting of ovarian cancer stem cells. Cancer Res, 2014. 74(17): p. 4922–36.

18. Wang, Y., et al., IL-6 mediates platinum-induced enrichment of ovarian cancer stem cells. JCI Insight, 2018. 3(23).

19. Chambers, C.R., et al., Overcoming the senescence-associated secretory phenotype (SASP): a complex mechanism of resistance in the treatment of cancer. Mol Oncol, 2021. 15(12): p. 3242–3255.

20. Nacarelli, T., et al., NAMPT Inhibition Suppresses Cancer Stem-like Cells Associated with Therapy-Induced Senescence in Ovarian Cancer. Cancer Res, 2020. 80(4): p. 890–900.

21. Hammen, P.K., et al., Multiple conformations of NAD and NADH when bound to human cytosolic and mitochondrial aldehyde dehydrogenase. Biochemistry, 2002. 41(22): p. 7156–68.

22. Parajuli, B., et al., Discovery of novel regulators of aldehyde dehydrogenase isoenzymes. Chem Biol Interact, 2011. 191(1-3): p. 153–8.

23. Parajuli, B., et al., Development of selective inhibitors for human aldehyde dehydrogenase 3A1 (ALDH3A1) for the enhancement of cyclophosphamide cytotoxicity. Chembiochem, 2014. 15(5): p. 701–12.

24. Buchman, C.D. and T.D. Hurley, Inhibition of the Aldehyde Dehydrogenase 1/2 Family by Psoralen and Coumarin Derivatives. J Med Chem, 2017. 60(6): p. 2439–2455.

25. Morgan, C.A. and T.D. Hurley, Development of a high-throughput in vitro assay to identify selective inhibitors for human ALDH1A1. Chem Biol Interact, 2015. 234: p. 29–37.

26. Tang, J., et al., Epigenetic Targeting of Adipocytes Inhibits High-Grade Serous Ovarian Cancer Cell Migration and Invasion. Mol Cancer Res, 2018. 16(8): p. 1226–1240.

27. Fang, F., et al., The novel, small-molecule DNA methylation inhibitor SGI-110 as an ovarian cancer chemosensitizer. Clin Cancer Res, 2014. 20(24): p. 6504–16.

28. Zong, X., et al., EZH2-Mediated Downregulation of the Tumor Suppressor DAB2IP Maintains Ovarian Cancer Stem Cells. Cancer Res, 2020. 80(20): p. 4371–4385.

29. Connor, E.V., et al., Thy-1 predicts poor prognosis and is associated with self-renewal in ovarian cancer. J Ovarian Res, 2019. 12(1): p. 112.

30. Wang, Y., et al., Frizzled-7 Identifies Platinum-Tolerant Ovarian Cancer Cells Susceptible to Ferroptosis. Cancer Res, 2021. 81(2): p. 384–399.

31. Ozes, A.R., et al., NF-kappaB-HOTAIR axis links DNA damage response, chemoresistance and cellular senescence in ovarian cancer. Oncogene, 2016. 35(41): p. 5350–5361.

32. Raspaglio, G., et al., Sox9 and Hif-2alpha regulate TUBB3 gene expression and affect ovarian cancer aggressiveness. Gene, 2014. 542(2): p. 173–81.

33. Li, J., et al., Lipid Desaturation Is a Metabolic Marker and Therapeutic Target of Ovarian Cancer Stem Cells. Cell Stem Cell, 2017. 20(3): p. 303–314 e5.

34. Hirschmann-Jax, C., et al., A distinct “side population” of cells with high drug efflux capacity in human tumor cells. Proc Natl Acad Sci U S A, 2004. 101(39): p. 14228–33.

35. Campisi, J. and F. d’Adda di Fagagna, Cellular senescence: when bad things happen to good cells. Nat Rev Mol Cell Biol, 2007. 8(9): p. 729–40.

36. Tomita, H., et al., Aldehyde dehydrogenase 1A1 in stem cells and cancer. Oncotarget, 2016. 7(10): p. 11018–32.

37. Meng, E., et al., ALDH1A1 maintains ovarian cancer stem cell-like properties by altered regulation of cell cycle checkpoint and DNA repair network signaling. PLoS One, 2014. 9(9): p. e107142.

38. Chefetz, I., et al., A Pan-ALDH1A Inhibitor Induces Necroptosis in Ovarian Cancer Stem-like Cells. Cell Rep, 2019. 26(11): p. 3061–3075 e6.

39. Levi, B.P., et al., Aldehyde dehydrogenase 1a1 is dispensable for stem cell function in the mouse hematopoietic and nervous systems. Blood, 2009. 113(8): p. 1670–80.

40. Feng, T., et al., Generation of mucosal dendritic cells from bone marrow reveals a critical role of retinoic acid. J Immunol, 2010. 185(10): p. 5915–25.

41. Cui, T., et al., DDB2 represses ovarian cancer cell dedifferentiation by suppressing ALDH1A1. Cell Death Dis, 2018. 9(5): p. 561.

42. Yokoyama, Y., et al., BET Inhibitors Suppress ALDH Activity by Targeting ALDH1A1 Super-Enhancer in Ovarian Cancer. Cancer Res, 2016. 76(21): p. 6320–6330.

43. Ozes, A.R., et al., Therapeutic targeting using tumor specific peptides inhibits long non-coding RNA HOTAIR activity in ovarian and breast cancer. Sci Rep, 2017. 7(1): p. 894.

44. Jordan, K.R., et al., The Capacity of the Ovarian Cancer Tumor Microenvironment to Integrate Inflammation Signaling Conveys a Shorter Disease-free Interval. Clin Cancer Res, 2020. 26(23): p. 6362–6373.

45. Gonzalez-Torres, C., et al., NF-kappaB Participates in the Stem Cell Phenotype of Ovarian Cancer Cells. Arch Med Res, 2017. 48(4): p. 343–351.

46. Uruski, P., et al., Primary high-grade serous ovarian cancer cells are sensitive to senescence induced by carboplatin and paclitaxel in vitro. Cell Mol Biol Lett, 2021. 26(1): p. 44.

47. Saleh, T., et al., Therapy-Induced Senescence: An “Old” Friend Becomes the Enemy. Cancers (Basel), 2020. 12(4).

