## Supplemental Data and Methods for "A novel ALDH1A1 inhibitor blocks platinum-induced senescence and stemness in ovarian cancer"

**Supplementary Figures**

**Supplementary Figure S1**

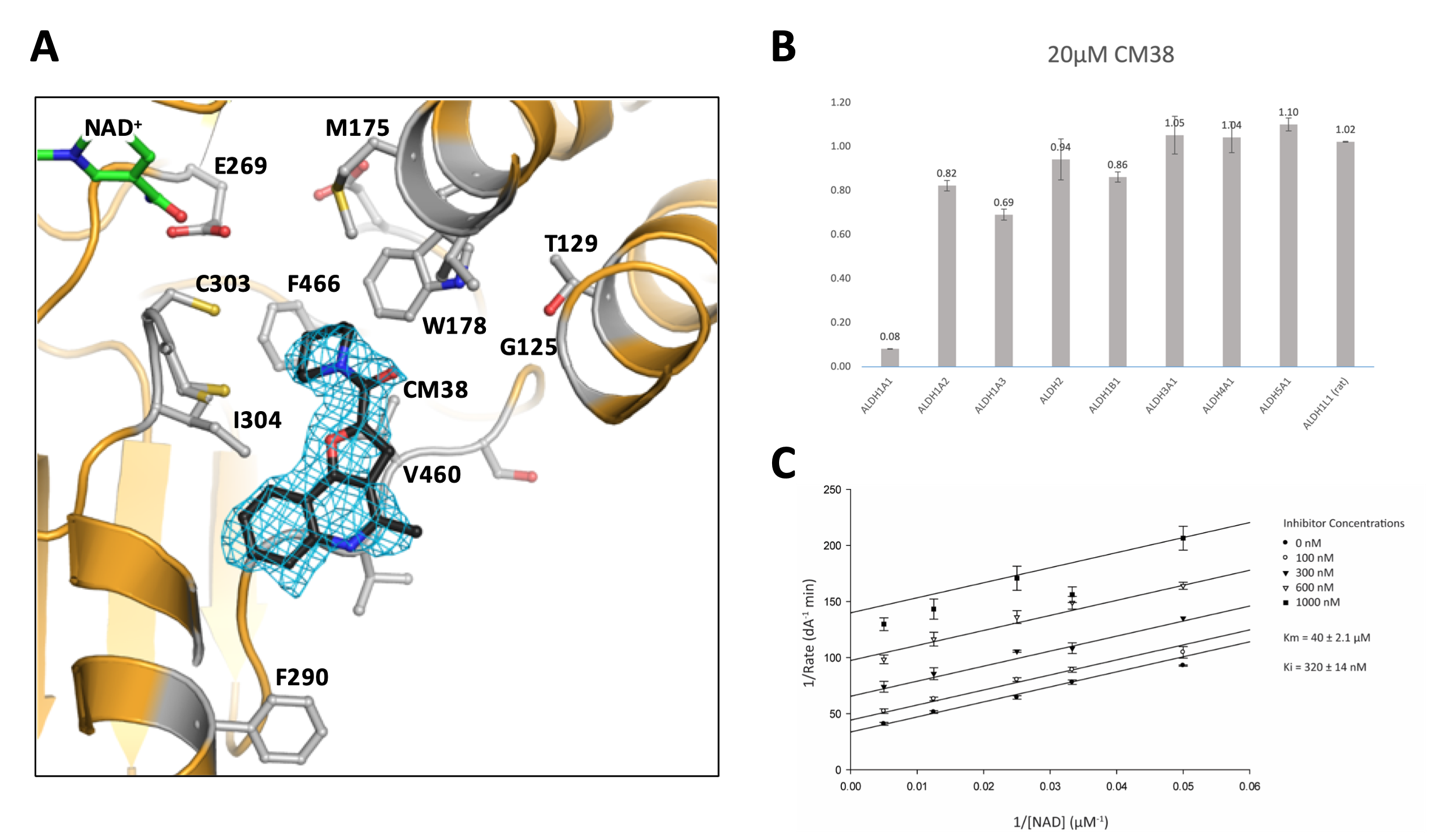

**Supplementary Figure S1**: X-Ray Crystallography of CM38 with ALDH1A1 and characterization of inhibitor activity **A.** Structure of Human ALDH1A1 with bound compound CM38. The figure shows the position of CM38 bound within the substrate binding pocket of human ALDH1A1 along with the observed electron density in the initial maps (m2Fo-DFc contoured at 1 standard deviation of the map) prior to inclusion of the ligand in the structure factor calculations. **B.** Inhibition of ALDH activity by compounds and IC50 curves were determined by measuring the formation of NAD(P)H spectrophotometrically at 340 nm using purified recombinant enzymes. **C.** Lineweaver-Burk Plot of CM38 Inhibition of ALDH1A1 with Varied NAD+ CM38 inhibition is uncompetitive with respect to varied NAD+ and has a Ki of 320 ± 14 nM.

**Supplementary Fig. S2**

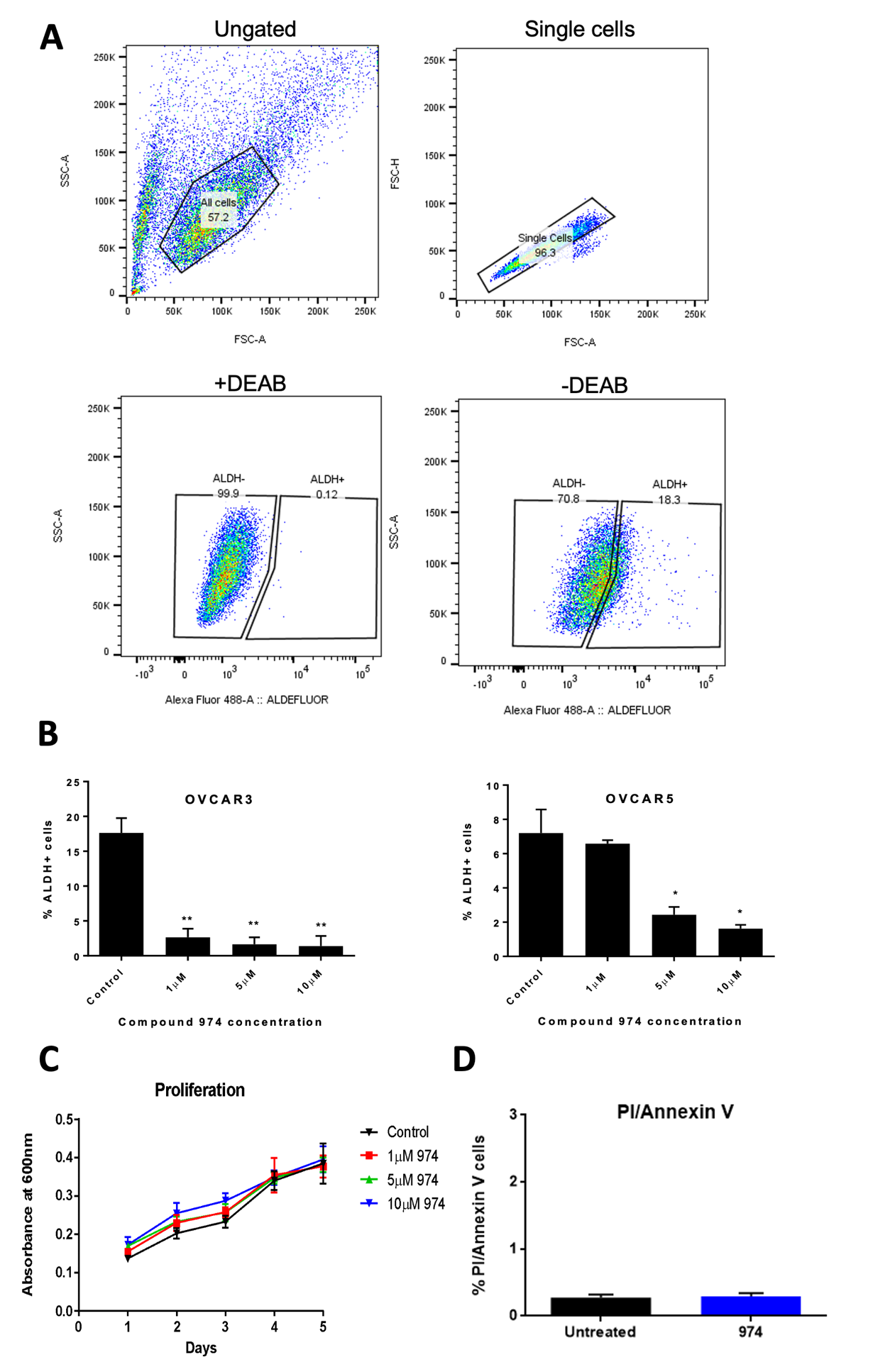

**Supplementary Fig. S2: Gating strategy for Aldefluor assay, dose response for 974, Proliferation and Apoptosis assay data A**. Gating strategy for ALDEFLUOR assay by flow cytometry in OVCAR3 cells. **B.** OVCAR3 (left) or OVCAR5 cells (right) were treated with 1,5 or 10 µM of 974 for 48h and ALDEFLUOR assay was performed. **C.** OVCAR3 cells were treated as in B and treated cells were plated at a density of 2000 cells/well and an MTT assay was performed on Day 1-5. **D.** OVCAR3 cells were treated with DMSO or 974 (5µM for 48h) and stained for PI/Annexin V and analyzed by flow cytometry.

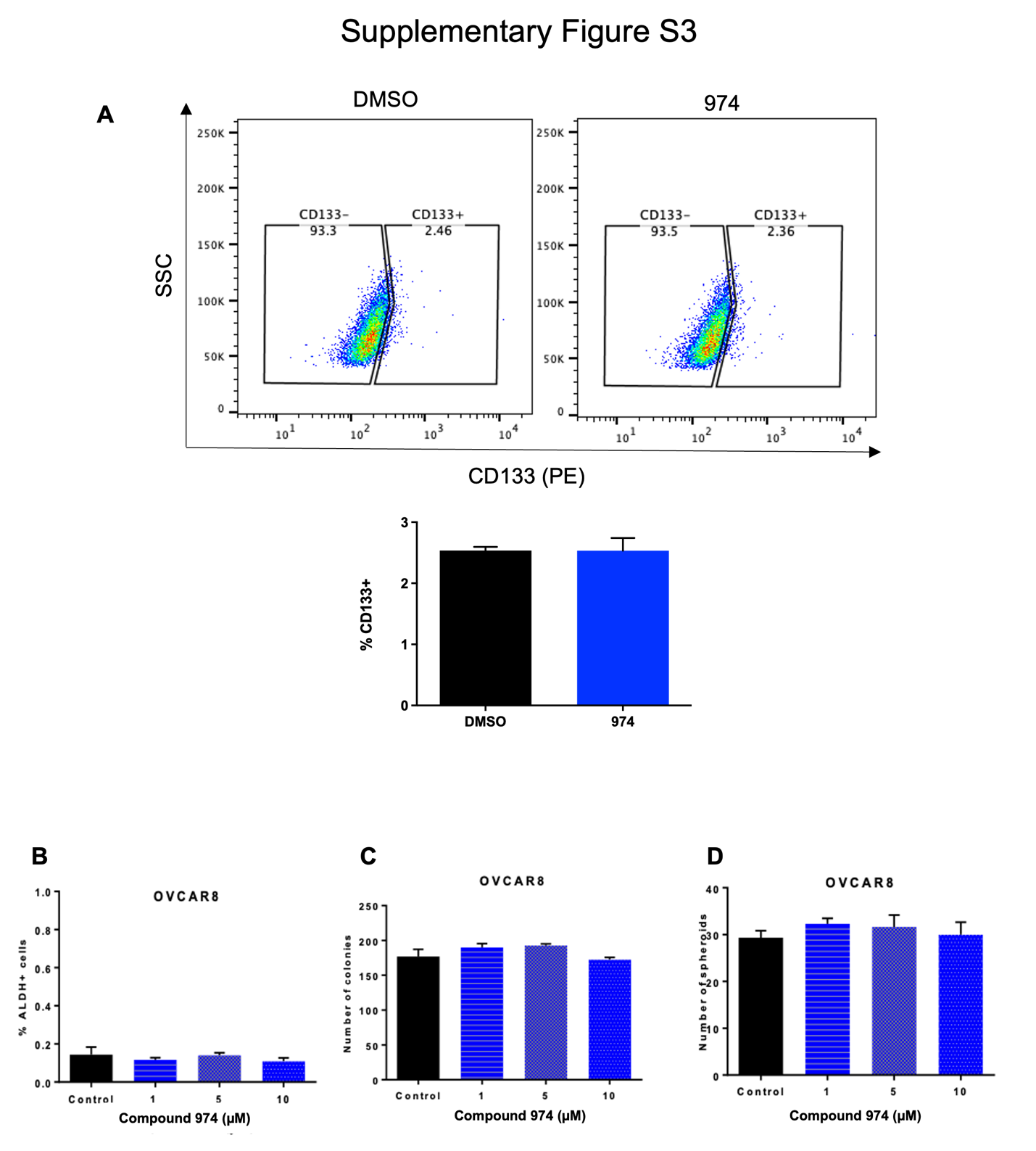

**Supplementary Fig. S3: CD133 expression and stemness assay validation in OVCAR8** OVCAR8 cells were treated with 1,5 or 10µM of 974 for 48h and

**A.** ALDEFLUOR assay was performed. **B.** 500 cells/well were replated in 6 well plates after treatment. Colonies were stained with crystal violet and counted. **C.** 500 cells/well were replated in 24-well non-adherent conditions after treatment. Spheroids were counted and quantified.

**D.** OVCAR8 cells treated with 5µM of 974 for 48h were stained for CD133 and analyzed for %CD133 cells by flow cytometry.

**Figure S4**

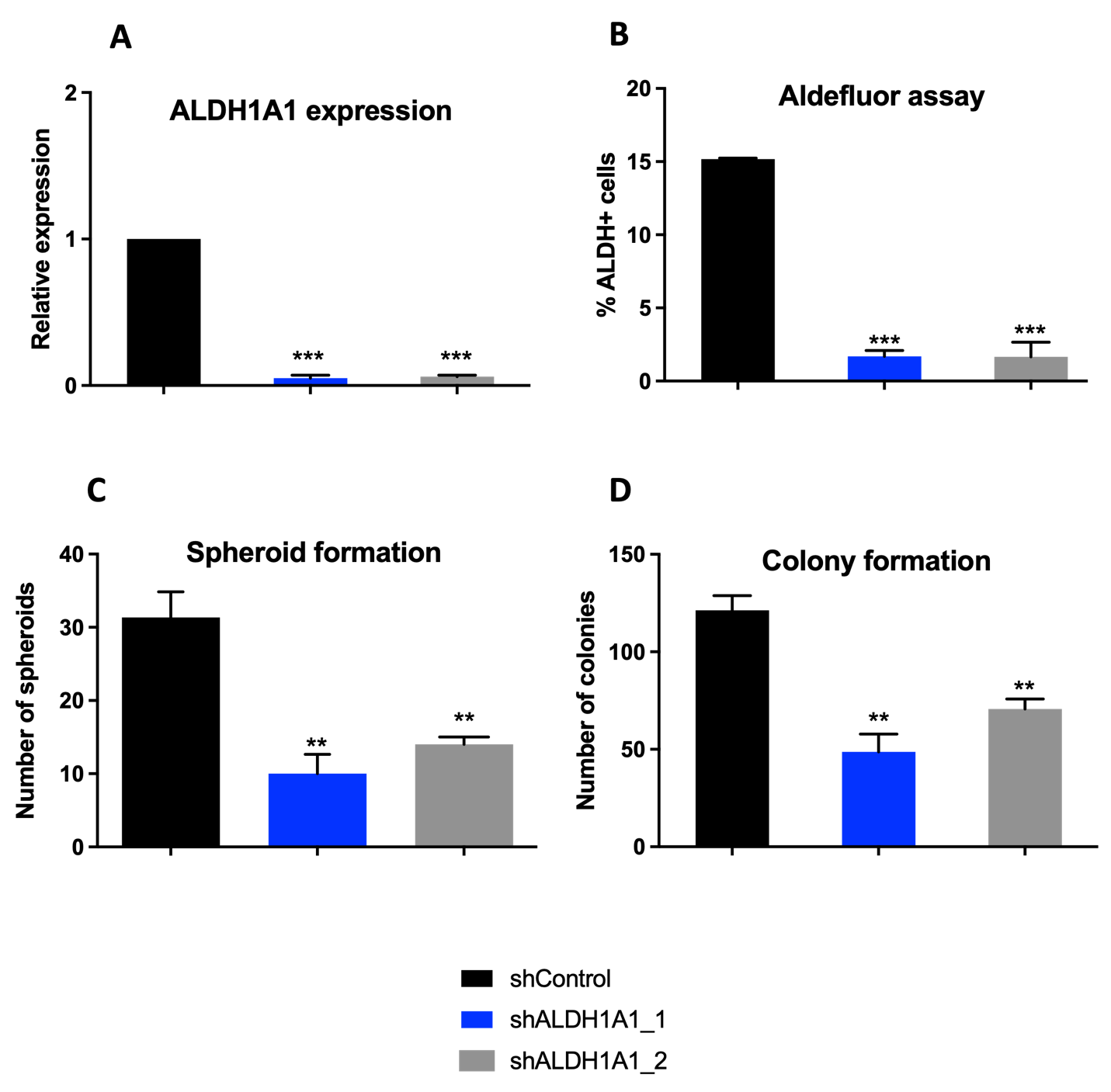

**Supplementary Figure S4: ALDH1A1 expression and stemness assays in ALDH1A1 shRNA knockdown and scrambled control cell lines.**

**A.** Expression of ALDH1A1 was measured by qPCR. **B**. % of ALDH+ cells in shControl or ALDH1A1 knockdown (shALDH1A1_1 or shALDH1A1_2) cells were measured by Aldefluor assay using flow cytometry. C. shControl or ALDH1A1 knockdown (shALDH1A1_1 or shALDH1A1_2) cells were plated at 500 cells/well were replated in 24-well non-adherent conditions. Spheroid formation was quantified after 14 days. **D.** shControl or ALDH1A1 knockdown (shALDH1A1_1 or shALDH1A1_2) cells were plated at 500 cells/well in 6 well plates. Colonies were stained with crystal violet and counted. Colony formation was quantified after 14 days. Error bars represent SEM; n =3 independent experiments of triplicate assays. Data are presented as mean ± SEM with p < 0.05 (*), p < 0.01 (**), and p < 0.005 (***).

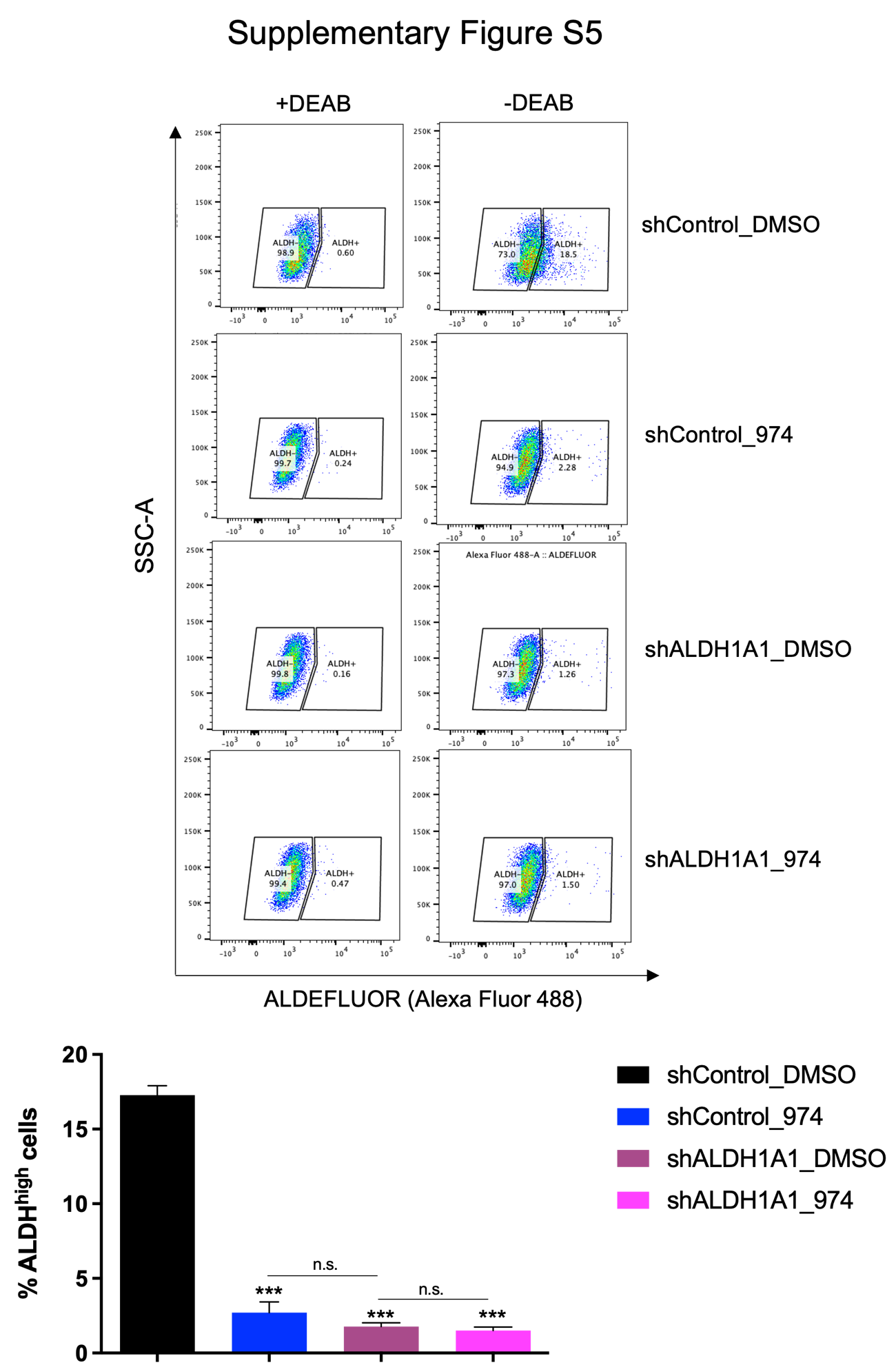

**Supplementary Figure S5: Aldefluor assay validation for ALDH1A1 knockdown and scrambled control cells treated with 974.**

**A.** shALDH1A1_1 or shControl cells were treated with DMSO or 974 and ALDEFLUOR assay was performed (top). % ALDH+ cells were quantified(bottom). Error bars represent SEM; n =3 independent experiments of triplicate assays. Data are presented as mean ± SEM with p < 0.05 (*), p < 0.01 (**), and p < 0.005 (***).

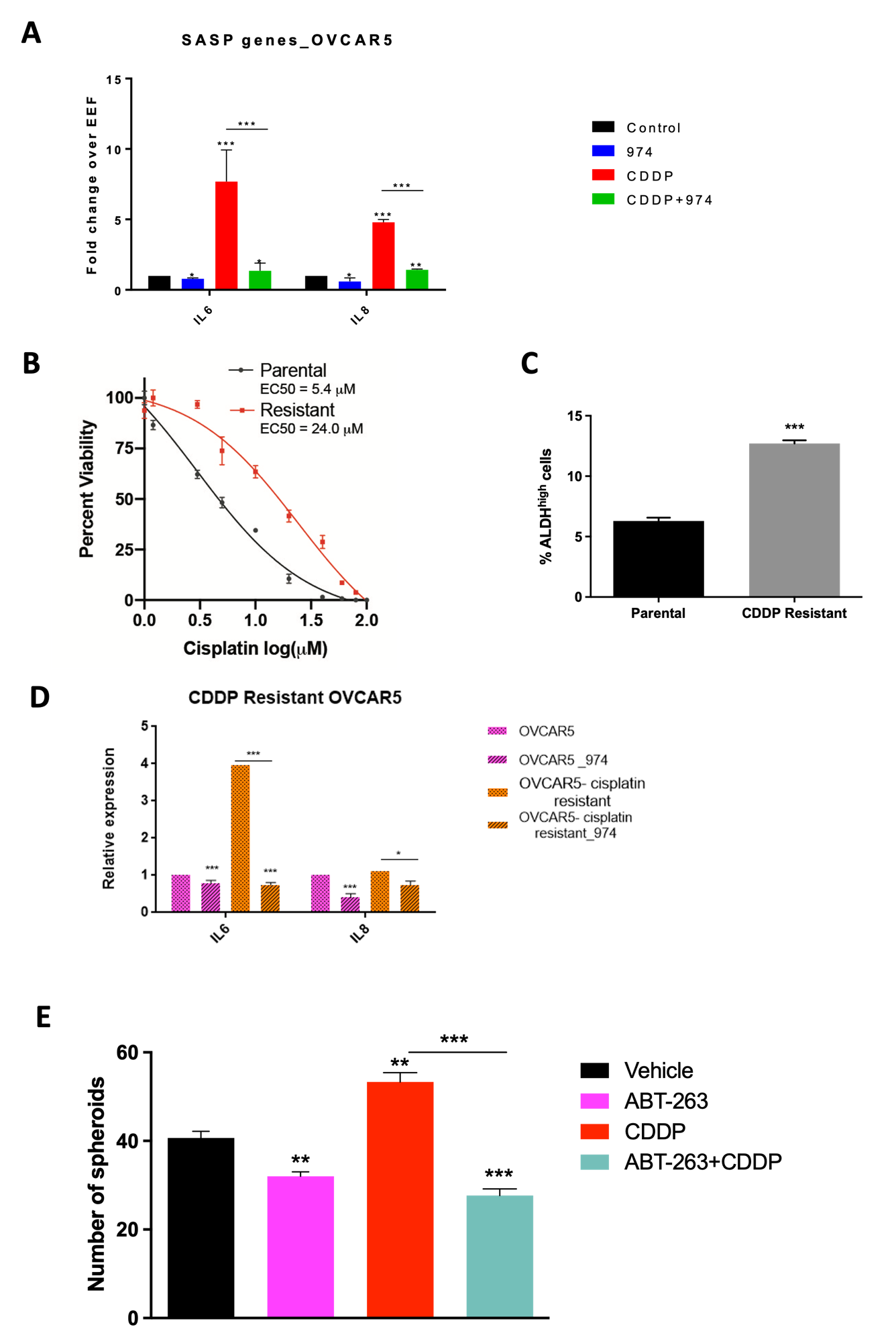

**Supplementary Figure S6: SASP gene expression in OVCAR5, characterization of CDDP-resistant OVCAR5 cells and SASP expression, spheroid formation in CDDP, ABT-263 combination**

**A.** Expression of SASP genes was examined by qPCR in OVCAR5 cells treated with DMSO, compound 974 (5µM for 48h), cisplatin (CDDP) (15µM for 16h) or both. **B.** EC50 curves for OVCAR5 parental and CDDP resistant lines generated by treating cells with an increasing dose of CDDP and MTT assay was performed to measure viability. EC50 was calculated using GraphPad Prism. **C.** ALDH activity was measured in OVCAR5 parental or CDDP resistant cells by ALDEFLUOR assay. **D.** Expression of SASP genes was examined by qPCR in parental or CDDP resistant OVCAR5 cells treated with DMSO or compound 974 (5µM for 48h). **E.** OVCAR3 cells were treated with ABT-263 (2µM for 24h) or), cisplatin (CDDP) (15µM for 16h) or both. 500 cells/well were replated in 24-well non-adherent conditions after treatment. Spheroid formation was quantified after 7 days.

**Supplementary Tables**

**Supplementary Table S1:** CM38 X-Ray Crystallography data

|  | **CM38** |
| --- | --- |
| PDB Code | 7UM9 |
| **Data Collection (ADSC Quantum Q210)** |  |
| Date of Collection  (Beamline 19-BM; wavelength 0.98 Angstroms) | 03/2016  19-ID |
| Space Group | P422 |
| **Cell Dimensions** |  |
| a,b,c (Å) | 109,109,83 |
| α,β,γ (deg) | 90,90,90 |
| Resolution (Å) | 50.0-1.80 |
| R_merge_ | 0.093 (0.50) |
| R_meas_ | 0.096 (0.53) |
| R_pim_ | 0.026 (0.16) |
| CC_1/2_ (highest shell) | 0.926 |
| I/σ_<I>_ | 24.4 (3.2) |
| Completeness (%) | 95.6 (77.6) |
| Redundancy | 12.9 (9.9) |
| **Refinement** |  |
| No. of reflections (R_w_) | 44,700 |
| No. of reflections (R_f_) | 2000 |
| No. of protein atoms | 3875 |
| No. of water molecules | 214 |
| No. of inhibitor molecules | 1 |
| Occupancy of inhibitor(s) | 1.0 |
| R_work_/R_free_ | 17.8/21.6 |
| **RMSD from Ideal Geometry** |  |
| Bond Length (Å) | 0.006 |
| Bond Angle (deg) | 0.90 |
| **Ramachandran plot** |  |
| Preferred (%) | 97.8 |
| Outliers (%) | 0.2 |
| Clashscore | 3.2 |
| Average B (Å) | 29.5 |
| Protein | 29.3 |
| Inhibitor | 35.7 |
| Solvent | 29.8 |

**Supplementary Table S2:** Primer sequences used for qPCR.

| **Primer Sequences** | **Forward** | **Reverse** |
| --- | --- | --- |
| EEF1A1 | TTGTCGTCATTGGACACGTAG | GATACCACGTTCACGCTCAG |
| Sox2 | GCGCGGGCGTGAACCAG | CGGCGCGGGGAGATACA |
| Oct4 | GGAAAGGCTTCCCCCTCAGGGAAAGG | AAGAACATGTGTAAGCTGCGGCCC |
| Bmi1 | TTACCTGGAGACCAGCAAGT | CATTAGAGCCATTGGCAGCA |
| Nanog | CAGAAGGCCTCAGCACCTACCTACCC | AAACCGACCTTGACGTACGTCCTGAC |
| IL6 | AACCTGAACCTTCCAAAGATGG | TCTGGCTTGTTCCTCACTACT |
| IL8 | GCTCTGTGTGAAG GTGCAGT | TGCACCCAGTTTTCCTTGGG |
| CXCL1 | AGTGGCACTGCTGCTCCT | TGGATGTTCTTGGGGTGAAT |
| CXCL3 | TCCGTGGTCACTGAACTGCG | AGTTGGTGCTCCCCTTGTTCA |
| CDKN1A (p21) | GGATGTCCGTCAGAACCC | GCTCCCAGGCGAAGTCA |
| ALDH1A1 | ATGGATGCTTCCGAGAGG | GCCACATACACCAATAGGTTC |
| ALDH1A2 | TTGGTTCAGTGTGGAGAAGG | AAAGCTTGCAGGAATGGTTTG |
| ALDH1A3 | CTTCTGCCTTAGAGTCTGGAAC | CGTATTCACCTAGTTCTCTGCC |
| ALDH3A1 | TGTTCTCCAGCAACGACAAG | CTGACCTTCAGGCCTTCATC |
| ALDH3A2 | ATTGTGGCTGT GGGTTGAGG | GAGGCACTAGGAGGTTGAA CAGG |

**Supplementary Table S3:** Compound activity against ALDH isoforms

| Compound | ALDH1A1  IC_50_ or % activity vs. control at 20 uM | ALDH1A2  IC_50_ or % activity vs. control at 20 uM | ALDH1A3  IC_50_ or % activity versus control at 20 uM | ALDH2  IC_50_ or % activity versus control at 20 uM | ALDH1B1  IC_50_ or % activity versus control at 20 uM |
| --- | --- | --- | --- | --- | --- |
| CM38  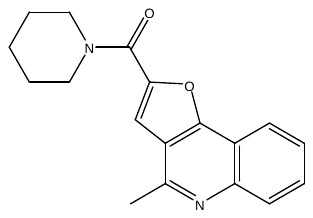 | 440 ± 37 nM | 82 ± 2.4 % | 69 ± 2.5% | 94 ± 9.4% | 86 ± 2.3% |
| 9040591  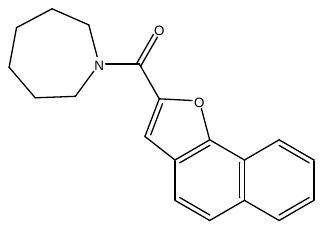 | 1500 ± 120 nM | 85 ± 1.3% | 103 ± 9.8% | 62 ± 1.0% | 94 ± 0.5% |
| C893-0789  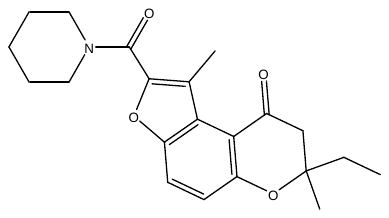 | 300 ± 14 nM | 58 ± 3.1% | 110 ± 3.2% | 85 ± 0.9% | 60 ± 1.9% |
| E003-0974  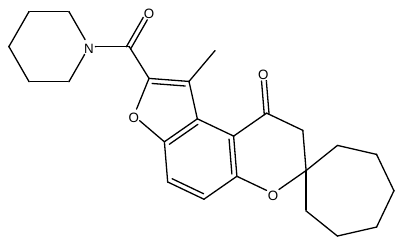 | 470 ± 29 nM | 63 ± 4.2% | 85 ± 4.0% | 76 ± 20% | 67 ± 3.9% |
| E003-0027  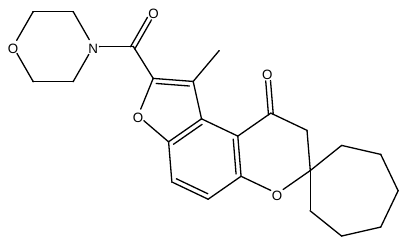 | 52 ± 2.1% | 47 ± 3.0% | 41 ± 0.8% | 64 ± 1.8% | 83 ± 0.9% |

**Supplementary Methods**

**Discovery of 974**

Compounds with high structural similarity to CM38 were ordered from commercial sources and evaluated as ALDH1A1 inhibitors. Variations that significantly decreased potency included removal of the piperidine ring, as well as lengthening the linkage between the aromatic core structure and the piperidine ring. Expansion of the piperidine ring to an azepane was tolerated as was the conversion of the pyridine ring to a benzene ring (9040591, **Supplementary Table S3**). However, removal of the methyl group from the central ring structure decreased selectivity toward ALDH2. 9040591 was used as a base to uncover additional analogs with central benzene ring structures. Due to the potential for reactivity due to the conjugated double bond system between the furan ring and the carbonyl group in 9040591, analogs were limited to those with methyl groups on the furan ring to impose steric restraints on attacking nucleophiles. Amongst these compounds, reducing the size of the piperidine ring to a pyrrolidine decreased potency significantly. Similarly, substitution to a morpholine group strongly reduces ALDH inhibition (E003-0027, **Supplementary Table S3**). The conversion of one of the aromatic rings in the structures to a dihydropyran-4-one (E003-0974, C893-0789, **Supplementary Table S3**) improves the potency of the compound toward ALDH1A1, improving the IC50 from 1.5 µM for 9040591 to 300 nM for C893-0789 or 470 nM for E003-0974. Of the two compounds, E003-974 demonstrated better activity in cells and was selected for bulk synthesis and further testing in cellular models.

**Bioinformatic analysis for RNA-seq**

Sequenced reads were adapter trimmed and quality filtered using Trimmomatic ver. 0.38 [1] setting the cutoff threshold for average base quality score at 20 over a window of 3 bases. Reads shorter than 20 bases post-trimming were excluded (parameters: LEADING:20 TRAILING:20 SLIDINGWINDOW: 3:20 MINLEN:20). Cleaned reads were mapped to Human genome reference sequence GRCh38.p12 with gencode v.28 annotation, using STAR, an RNA-seq aligner [2] version STAR_2.7.3a (parameters: --outFilterMultimapNmax 20 --seedSearchStartLmax 16 --twopassMode Basic). Gene expression was quantified by counting the number of reads mapped to the exon regions of the annotated genes using featureCounts tool ver. 2.0.0 of subread package [3](parameters: -s 2 -p -B -C). Differential Expression Analysis was performed using DESeq2 ver. 1.12.3[4]. The p-values for genes indicating the probabilities of no differential expression across the sample groups were corrected for multiple testing using Benjamini-Hochberg method.

**Generation of cisplatin resistant OVCAR5 cells**

To generate cisplatin resistant cells, OVCAR5 cells were treated with IC70 dose of cisplatin (18 μM) for 3 hours. Following treatment, media was removed, cells were washed two times with PBS, fresh media without cisplatin was added back to the flask and the cells were allowed to recover. Once cells reached 60-70% confluency, cells were split at 1:2. When cells reached 80% confluency the cisplatin treatment was repeated as the second cycle of treatment. Cells received 5 cycles of treatment and a maintenance treatment with 18 μM cisplatin was performed every month to retain a resistant population.

1.	Bolger, A.M., M. Lohse, and B. Usadel, *Trimmomatic: a flexible trimmer for Illumina sequence data.* Bioinformatics, 2014. **30**(15): p. 2114-20.

2.	Dobin, A., et al., *STAR: ultrafast universal RNA-seq aligner.* Bioinformatics, 2013. **29**(1): p. 15-21.

3.	Liao, Y., G.K. Smyth, and W. Shi, *featureCounts: an efficient general purpose program for assigning sequence reads to genomic features.* Bioinformatics, 2014. **30**(7): p. 923-30.

4.	Love, M.I., W. Huber, and S. Anders, *Moderated estimation of fold change and dispersion for RNA-seq data with DESeq2.* Genome Biol, 2014. **15**(12): p. 550.
